# Evaluation of Metrics for Benchmarking Antimicrobial Use in the United Kingdom Dairy Industry

**DOI:** 10.1101/186593

**Authors:** Harriet L. Mills, Andrea Turner, Lisa Morgans, Jonathan Massey, Hannah Schubert, Gwen Rees, Fraser Broadfoot, David Barrett, Andrew Dowsey, Kristen K. Reyher

## Abstract

The issue of antimicrobial resistance is of global concern across human and animal health. In 2016 the UK government committed to new targets for reducing antimicrobial use (AMU) in livestock. However, though a number of metrics for quantifying AMU are defined in the literature, all give slightly different interpretations.

This paper reviews a selection of metrics for AMU in the dairy industry: total mg, total mg/kg, daily dose and daily course metrics. Although the focus is on their application to the dairy industry, the metrics and issues discussed are relevant across livestock sectors.

In order to be used widely, a metric should be understandable and relevant to the veterinarians and farmers who are prescribing and using antimicrobials. This means that clear methods, assumptions (and possible biases), standardised values and exceptions should be published for all metrics. Particularly relevant are assumptions around the number and weight of cattle at risk of treatment and definitions of dose rates and course lengths; incorrect assumptions can mean metrics over- or under-represent AMU.

The authors recommend that the UK dairy industry work towards UK-specific metrics using UK-specific medicine dose and course regimens as well as cattle weights in order to monitor trends nationally.

## Introduction

Antimicrobial resistance (AMR) is a matter of global concern across the human health, animal health and agricultural sectors. In May 2015, the World Health Organisation (WHO) published a Global Action Plan to tackle AMR (1) which identified the need for strong collaborations between the three sectors to address the problem. The WHO also publish and maintain a list of antimicrobials (AMs) which are of critical importance to human health (2). This list of Critically Important Antimicrobials (CIAs) has been further refined by the European Medicines Agency (EMA) to give a list of Highest Priority CIAs (HP-CIAs: fluoroquinolones, 3^rd^ and 4^th^ generation cephalosporins and colistin (3, 4)) and these have been adopted by the UK (5). Globally, it is recognised that the use of HP-CIAs across sectors needs to be reduced, and the EMA has called for significant restrictions in their use in animals (6).

In 2014 the UK government commissioned a review to analyse the problem of AMR globally and propose solutions. This review (7) (published mid-2016) specifically called for a reduction in the use of AMs in farming and an increase in the regulatory oversight of antimicrobial use (AMU) and resistance in animals. The UK government response (published September 2016) committed to decreasing overall average AMU in livestock to 50 mg/kg by 2018, a 19% reduction from 62 mg/kg in 2014 (8). This 50 mg/kg target is for the livestock industry as a whole. AMU, however, varies considerably across livestock sectors and therefore the government has also committed to working with individual sectors to set appropriate sector-specific reduction targets by 2017. These targets will focus on encouraging best practice and responsible use of antimicrobials, as well as safeguarding animal health and welfare.

There has been a concerted effort in the livestock industry to raise awareness of AMU, with some food retailers and milk buyers placing emphasis on regular reporting of usage data. A variety of metrics (measures) for AMU are used across the industry and have been presented and used in the literature (5, 9-17). All of these seek to monitor changes in AMU over time, assess the impact of policy change, and, potentially, benchmark farms or veterinarians against one another. However, each metric comes with its own assumptions, meaning that each gives a slightly different interpretation and view of AMU. In the opinion of these authors, an ideal metric needs to be easily comparable across different units (farms, veterinarians, regions), and take into account the number and range of animals (ages, breeds, etc.) to which AMs are being prescribed. In order to be used widely in the livestock industries, such a metric should also be understandable and relevant to veterinarians and farmers who are prescribing and using AMs. Clear methods, assumptions (and possible biases), standardised values and exceptions should be published for all metrics such that each may be independently calculated and compared, and the overriding goal of reducing AMR should be kept in mind at all times.

The aim of this manuscript is to specifically focus on the pros and cons of a selection of metrics for measuring AMU in the UK dairy cattle industry, although most metrics presented can be applied to other types of livestock.

## Metrics

Five metrics for AMU are described in the following section.

### Total mg

**Total mg** of active substance is simple to calculate and easy to understand. However, it ignores variation in dose rates across AMs and individual differences between farms and veterinarians. For example, one farm may compare favourably to another only because of dose rate differences in the medicines they use; importantly, this is especially true for the HP-CIAs, which tend to have low dose rates (Table 1). Total mg is also not suitable for comparison across farms with different numbers of cattle: farms using the same amount of a particular medicine per animal will have different total mg depending on the number of cattle on each operation. On farms with smaller or lighter animals, total mg will be lower even if the number of doses per animal is the same as a farm with larger or heavier animals. For cattle, AMs (such as lincomycin and tylosin) are sometimes used in footbaths in a way that does not follow the clinical recommendations on the Summary of Product Characteristics (SPC), under the Cascade system (18). This use could be at such quantities that the increase in total mg for active substances in those AMs would be heavily inflated compared to farms not using AM footbaths - this applies to all AMU metrics.

**Table 1:** Demonstrating the different total mg and total mg/kg between two (hypothetical) farms with the same total kg of treated animals who are using AMs with different dose rates. Farm 1 is using a 3^rd^ generation cephalosporin (an HP-CIA), but, as the dose rate of cephalosporins is lower than that of tetracyclines, the total mg of medicine used on Farm 1 is less than on Farm 2, so both metrics appear to be lower for Farm 1 than Farm 2. The metrics ignore the fact that Farm 1 is using an HP-CIA, which is arguably more important to reduce than just minimising total use when considering selection for antimicrobial resistance.

| Farm | Medicine | AM type | Dose rate (mg/kg) | Concentration (mg/ml) | Total weight of treated animals (kg) | AMU |  |
| --- | --- | --- | --- | --- | --- | --- | --- |
|  |  |  |  |  |  | Total mg | Total mg/kg |
| 1 | Ceftiofur (Naxcel™, Zoetis, UK) | 3rd generation cephalosporin | 6.6 | 200 | 60,000 | 396,000 | 6.6 |
| 2 | Oxytetracycline (Terramycin LA™, Zoetis, UK - long-acting dose) | Tetracycline | 20 | 200 | 60,000 | 1,200,000 | 20 |

### Total mg/kg

**Total mg/kg** (5) improves on total mg by dividing the mass of the medicines by the total weight of cattle at risk of treatment, therefore accounting for variation in cattle numbers and weights across farms. However, as with total mg, use of this metric may encourage favouring of the HP-CIAs for their lower mg per dose (Table 1). O’Neill’s AMR Review recommended a reduction in the use of the HP-CIAs, although they did not specifically suggest a separate target (7). In order to prevent a shift towards the HP-CIAs to meet an overall mg/kg figure, there should always be a separate calculation for HP-CIAs (as is shown in the UK VARSS reports (5)). In the drive to reduce AMR, it is necessary to recognise that, in some instances, using more mg of medicine (moving from the use of fluoroquinolones to tetracyclines, for instance) may actually be beneficial.

Commonly, actual cattle weights on farms are not known and so most systems rely on estimated weights. The published literature presents a large range of cattle weights; for example, weights used for adult milking cattle range from 425 kg (estimated mean weight at time of treatment defined by the European Surveillance of Veterinary Consumption (ESVAC) group (19): if this weight is used, the metric is commonly referred to as **mg/PCU** (Population Correction Unit (17)), 600 kg (used by Netherlands and Denmark for national reporting (11, 13)) to 680 kg (20). Cattle weight also varies by age and breed, with the additional complication that many herds are of mixed breeds, making the use of a standard weight potentially problematic. Additionally, many AMs are specifically (or predominantly) used in youngstock, dairy or beef cattle and there is variation in disease susceptibility between breeds (21, 22). If an average cattle weight is known for the farm (through systems such as robotic milking machines that have a weigh floor), or if an average weight for the farm’s most common breed is used, and/or use is divided by age, metrics will give a more accurate result for the farm. Data to inform current mean weights of UK cattle for different breeds have been collected and these up-to-date estimates will help improve accuracy in UK metrics (H. Schubert, S. Wood, K.K. Reyher, H.L. Mills, in preparation).

Using an inaccurate weight for the animals at risk of treatment on a farm may result in any of the ‘per kg’ metrics under-or over-representing actual AM use, thereby rendering comparisons across farms with different mean weights (for example due to different breeds) inaccurate (Table 2).

**Table 2:** Demonstrating the problem of farm-specific cattle weights vs. standard weights when comparing AMU in total mg/kg across (hypothetical) farms. Both farms have the same number of cattle and use the same total mg of medicine, but cattle on Farm 3 are heavier than those on Farm 4 (e.g. Holstein vs. Jersey). Total mg/kg using a cattle weight specific to that farm gives more accurate figures than using a standard weight (in this case, the standard weight chosen is 425 kg, the estimated mean weight at time of treatment for dairy cattle defined by the European Surveillance of Veterinary Consumption group, ESVAC).

| Farm | Total mg | Total number of adult milking cattle | Mean cattle weight (kg) | AMU |  |
| --- | --- | --- | --- | --- | --- |
|  |  |  |  | Total mg/kg using mean cattle weight specific for that farm | Total mg/kg using ESVAC 425 kg mean weight at time of treatment |
| 3 | 1,000,000 | 100 | 750 | 13.3 | 23.5 |
| 4 | 1,000,000 | 100 | 450 | 22.2 | 23.5 |

‘Per kg’ metrics are also subject to further inaccuracies and lack of comparability between users if the total kgs of animal at risk of treatment take different animal populations into account. For example, if only adult milking cattle are included when calculating total kgs, a dairy farm that rears its own youngstock will have the same kg weight assigned as an equivalent farm that does not rear youngstock, even though there are more animals at risk of treatment with AMs (Table 3). Similarly, if all animals on the holding are included, a dairy farm keeping beef animals is likely to have a lower mg/kg when compared with a dairy-only farm with the same number of animals, due to the relatively low use of AMs in beef animals when compared to dairy.

**Table 3:** Demonstrating how the animal population included in the total kgs treated calculation can influence the final value for total mg/kg. Here, both (hypothetical) farms have the same number of adult cattle, but Farm 5 does not have youngstock; if only adult cattle are included in total kg then the total mg/kg are the same for both farms. If both adults and youngstock are included then total mg/kg is lower for Farm 6 (which does have youngstock).

| Farm | Total mg | Total number of adult milking cattle | Total weight of adult milking cattle using 600 kg standard weight (kg) | Total number of youngstock <12 months | Total weight youngstock <12 months using 100 kg standard weight (kg) | AMU |  |
| --- | --- | --- | --- | --- | --- | --- | --- |
|  |  |  |  |  |  | Total mg/kg (including only adult milking cattle weight) | Total mg/kg (including both adult milking cattle and youngstock weight) |
| 5 | 1,000,000 | 100 | 60,000 | 0 | 0 | 16.7 | 16.7 |
| 6 | 1,000,000 | 100 | 60,000 | 50 | 5,000 | 16.7 | 15.4 |

An alternative to mg/kg would use production data instead of weight, such as mg/1000 L of milk produced. These sorts of metrics might be valued by some farmers. There have, however, been suggestions that metrics taking into account production data imply to the public that AMs are present in animal products at substantial levels, which is misleading to consumers.

### Daily Dose metrics

**Defined Daily Dose (DDD) metrics** divide the total mg of medicine used by both total animal weight and an estimate of the daily dose for that medicine. These metrics are commonly used in human medicine (23) and help to overcome the issue of total mg and mg/kg metrics not accounting for different dose rates in AMs (highlighted in Table 1). As well as using either actual or standard weights for animals at risk of treatment (see mg/kg), daily dose metrics can use “*actual”* daily doses (e.g. farm-specific) or *“defined”* daily doses (e.g. recommended or standard doses; Figure 1).

**Figure 1:**
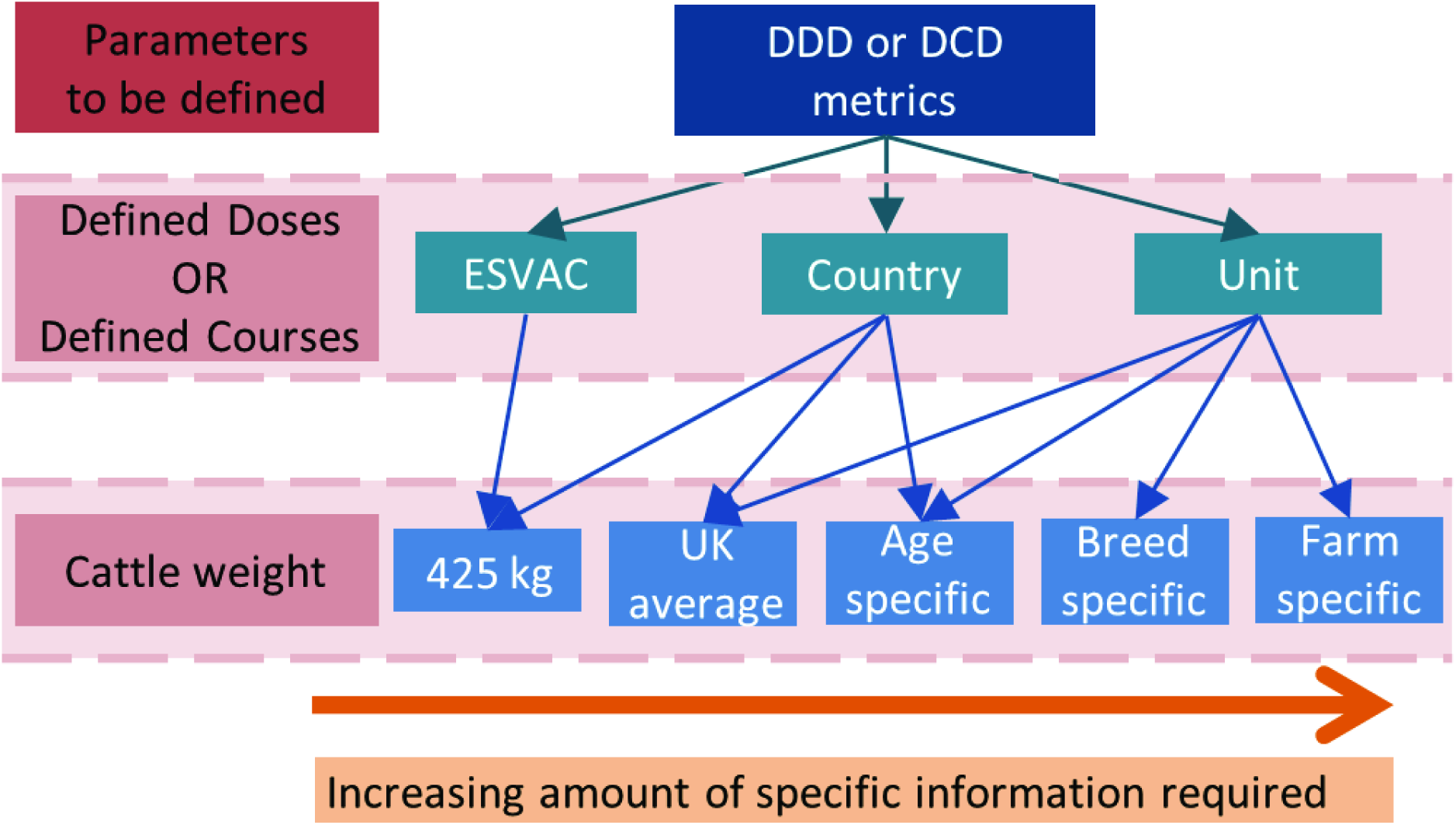
Flowchart explaining the different options for defined daily dose (DDD) and defined course dose (DCD) metrics. These metrics can (and have) been defined at European Surveillance of Veterinary Consumption (ESVAC) group, country or unit level, requiring doses or courses to be defined or known specifically for that level. These metrics also require cattle weights, and the ESVAC, country or unit level may define weight in different ways. The amount of specific information required increases from left to right in the figure as metrics become more representative of the actual situation on farms, with a tradeoff of increasingly granular information necessary to calculate the metrics.

The ESVAC group have formalised a **Defined Daily Dose for animals (DDDvet)** metric for dairy cattle which uses fixed daily dose definitions and a standard weight of 425 kg (estimated mean weight at time of treatment for dairy cattle) (10). Daily doses for DDDvet are defined per active substance and administration route rather than per individual product, and are based on the arithmetic mean dose of all veterinary medicine products, given by the standard product documentation from nine countries: Czech Republic, Denmark, Finland, France, Germany, the Netherlands, Spain, Sweden and the United Kingdom. Because these definitions represent an average across countries and do not take into account within-product variation they may not reflect actual prescription and use practices in an individual country, meaning DDDvet may be less representative at a country, farm or veterinary practice level.

Daily dose definitions have been published for all products except long-acting gamithromycin and tildipirosin (which will be published at a later date) (10). For dairy cattle specifically, there is a problem accounting for use of intramammary tubes. These have low mg per dose (and therefore do not substantially increase mg/kg), but do impact the number of daily doses administered. Currently, dry cow antibiotic tubes have not been assigned a DDDvet value, though lactating cow tubes have (1/teat) (10). Another issue for cattle is the inclusion of AMs used under the Cascade in footbaths - because there are no defined doses for this method, this use cannot be included in daily dose metrics. However, AMs can be used at very high quantities in footbaths, meaning that excluding them can under-represent actual AMU on farms.

To improve representativeness, daily dose metrics can be defined at country level (i.e. the fixed daily dose definitions and standardised weights would be specific to that country) or at the unit level (e.g. farms or veterinary practices, by using the individualised dose regimens and even weights actually reported by the farm or veterinary practice; Figure 1). These versions are potentially powerful, as the inclusion of more accurate data improves the representativeness of the metric and allows better comparisons across countries or units (15).

Whether the minimum, mean or maximum recommended rates are chosen as the defined dose rate significantly impacts the final DDD metric, illustrating how different choices - even taken within the recommended range - could alter the interpretation of AMU (Table 4). These biases also apply if the actual dose rate used on the farm is different to the defined dose rate: for example, the maximum dose rate may often be administered on farms so the mean may not accurately reflect use. The choice of animal weight can also cause similar biases, as previously discussed (Table 2).

**Table 4:** Demonstrating the impact different dose rates may have on the final DDD metric. This table uses tylosin as an example, showing the difference in DDD when taking the minimum, mean or maximum recommended dose rate (24). Tylosin has a range of dose rates, resulting in a range of DDD values.

| Medicine | Dose rate (mg/kg) | Total number of adult milking cattle | Total weight of adult milking cattle using 600 kg standard weight (kg) | AMU |  |  |
| --- | --- | --- | --- | --- | --- | --- |
|  |  |  |  | Total mg | Total mg/kg | Defined Daily Doses |
| Tylosin | 4 (minimum) | 100 | 60,000 | 100,000 | 1.67 | 0.42 |
| Tylosin | 7 (mean) | 100 | 60,000 | 100,000 | 1.67 | 0.24 |
| Tylosin | 10 (maximum) | 100 | 60,000 | 100,000 | 1.67 | 0.17 |

Note that for different countries and AMU monitoring systems, daily dose metrics have also been termed Animal Daily Dose (ADD), Defined Animal Daily Dose (DADD) and Defined Daily Dose Animal (DDDA). Calculations are the same, but different countries and systems use different daily doses and cattle weights, and include different specific (e.g. age) groups.

### Course Dose metrics

**Course dose metrics** attempt to assign the number of courses an animal receives, taking into account the daily dose and the course length. The ESVAC group have formalised a **Defined Course Dose for animals (DCDvet)** (10) as a suitable metric for monitoring across the EU. DCDvet is similar to DDDvet, but uses fixed <u>course dose</u> definitions instead of fixed <u>daily dose</u> definitions (based on the same nine European countries as DDDvet) as well as an assumed weight of 425 kg. These assumptions introduce the same problems as for DDDvet discussed above. Unlike DDDvet, however, both intramammary lactating and dry cow tubes have DCDvet values: 3/teat for lactating cow tubes and 4/udder for dry cow tubes (10).

As with daily dose metrics, if actual dosage regimens, course lengths and cattle weights are used, these would produce the most accurate DCD metric for each unit (Figure 1). However, this level of detail is not always available.

### Cow Calculated Course

**Cow Calculated Course (CCC)** is a metric conceived in the UK as part of an XLVet initiative (T. Clarke, personal communication). This metric uses course length data and dosing regimen as per the UK SPC documents and the number of cattle on the holding (taken from the Cattle Tracing System, which uses British Cattle Movement Systems (BCMS) data). CCC splits out medicine use into youngstock and adult stock by assuming certain products are only used in certain age groups. Udder preparations and short-acting injectable antibiotics are allocated to adults, and long-acting injectable and oral antibiotic products are deemed as youngstock treatments. CCC tallies the courses of each medicine used in a set time period and divides this by the number of animals on the holding. CCC makes assumptions on cattle weight (100 kg for youngstock (<24 months) and 600 kg for adult dairy animals (>24 months)) in order to work out how many courses are in a given saleable unit of medicine. When the course length is a range of days on the SPC, CCC uses the longest course length and the highest dose rate as assumptions for calculating how many courses one saleable unit of medicine contains. To make the metric more accurate at a farm level, the actual course length per medicine as given by the farmer and ideally the on-farm cattle weights and dose rates per medicine could be used (Figures 1 & 2). Although these parameters should be derivable from on-farm records, this level of detail may not be easy to collect.

**Figure 2:**
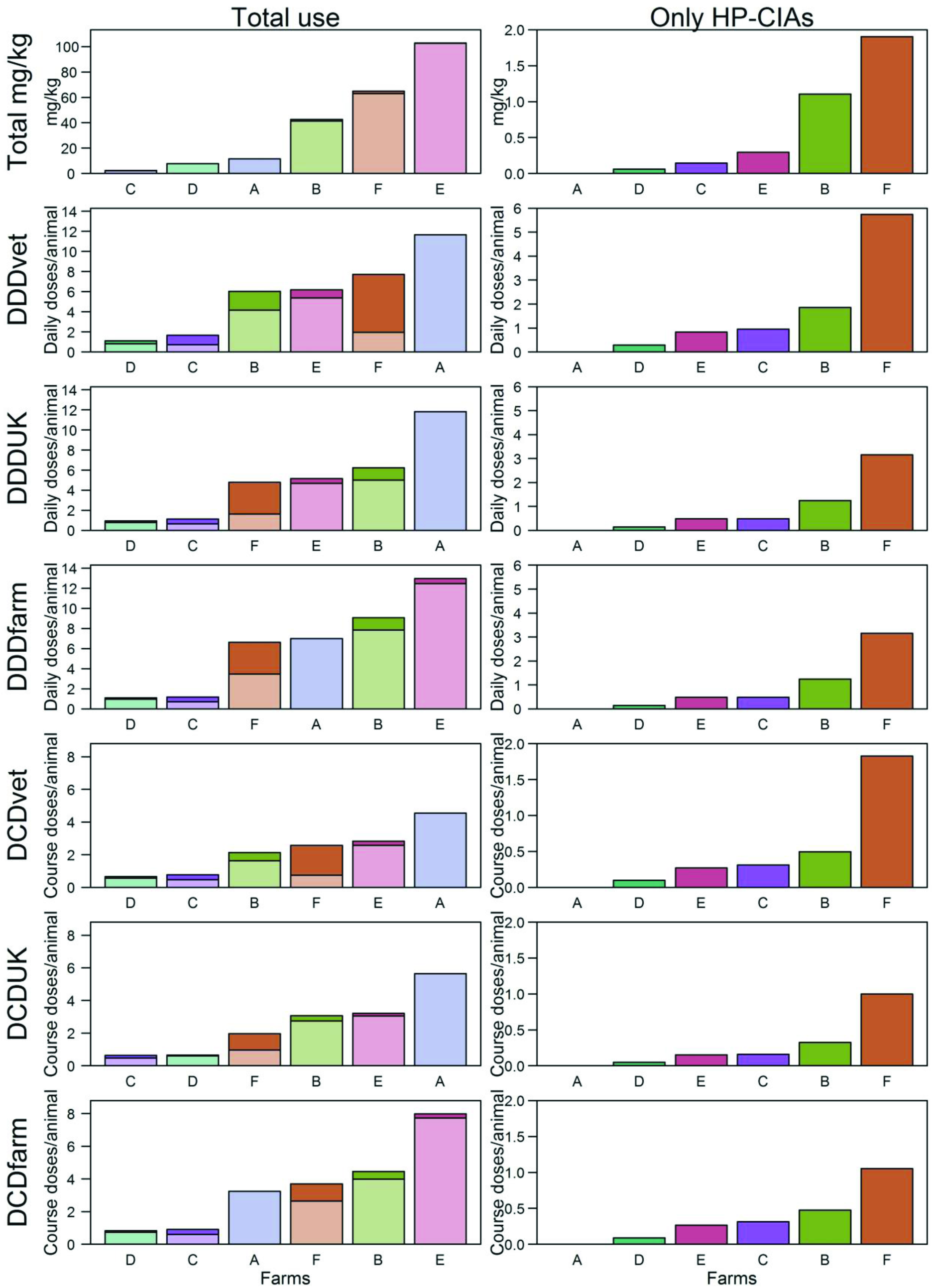
Antimicrobial use for six farms over a twelve-month period illustrated as total mg/kg, DDDvet, DDDUK, DDDfarm, DCDvet, DCDUK and DCDfarm. Figures on the left show total use with the darker shading indicating the highest priority critically important antimicrobials (HP-CIAs); figures on the right show HP-CIA use exclusively. Farms are coloured differently and plotted in order of their use for each metric.

**Table 5:** Definitions for different metrics for presenting antimicrobial use including calculations, data required and pros and cons for each.

| Metric | Calculation | Requirements | Pros | Cons (including assumptions) | Reference |
| --- | --- | --- | --- | --- | --- |
| Total mgs | = (Total mls, number or kg of product sold or used) x (mg/ml, mg/unit or mg/kg) | • Total mg of each active substance used | • Simple | • Ignores variation in animal size/weight and number of cattle<br>• Does not take into account different dose rates and course lengths across AMs | - |
| Total mg/kg<br>(sometimes called mg/PCU if the standard weight of 425 kg is used) | = Total use (mg) / total weight of cattle at risk of treatment (kg)<br>(total use (mg) defined as above) | • Total mg of each active substance used<br>• Number of cattle at risk of treatment<br>• Cattle weight (can be ESVAC defined (425 kg) or country- or unit-specific) | • Simple<br>• Takes into account animal weight and number of cattle | • If using standardised weights these may not be accurate for the unit in question<br>• Does not take into account different dose rates and course lengths across AMs | (5) |
| Daily Dose metrics<br>(e.g. DDDvet) | = Total use (mg) / (daily dose (mg/kg) x total weight of cattle at risk of treatment (kg))<br>The calculation is carried out for each AM category and added together to get the total figure.* | • Total mg of each active substance used<br>• Daily dose per kg* - these can be ESVAC-defined, or country- or unit-specific<br>• Number of cattle at risk of treatment<br>• Cattle weight (can be ESVAC-defined (425 kg) or country- or unit-specific) | • Takes into account animal weight and number of cattle<br>• Takes into account dose rate differences across AMs<br>• ESVAC-supported<br>• Dose and cattle weight can be made specific to country or unit which improves accuracy and comparability | • More complicated to calculate than mg and mg/kg metrics<br>• If using standardised weights and/or defined dose rates, these may not be accurate for a particular country or unit<br>• Assumptions rely upon knowledge of certain aspects of medicine pharmacokinetics that are not necessarily known (e.g. the exact duration of action) | (10) |
|  |  |  |  | • Does not take into account different course lengths across AMs<br>• Some drugs or routes of administration do not have defined dose values |  |
| Course Dose metrics (e.g. DCDvet) | = Total use (mg) / (course dose (mg/kg) x total weight of cattle at risk of treatment (kg))<br>The calculation is carried out for each antibiotic category and added together to get the total figure. | • Total mg of each active substance used<br>• Daily course per kg - these can be ESVAC-defined, or country- or unit- specific<br>• Number of cattle at risk of treatment<br>• Cattle weight (can be ESVAC-defined (425 kg) or country- or unit-specific) | • Takes into account animal weight and number of cattle<br>• Takes into account dose rate and course length differences across AMs<br>• ESVAC-supported<br>• Dose, course and cattle weight can be made specific to country or unit which improves accuracy and comparability | • More complicated to calculate than mg and mg/kg metrics<br>• If using standardised weights and/or defined dose rates and course lengths, these may not be accurate for a particular country or unit<br>• Assumptions rely upon knowledge of certain aspects of medicine pharmacokinetics that are not necessarily known (e.g. the exact duration of action) | (10) |
| Cow Calculated Courses (CCC) | = Number of courses in 12 months / number of cattle on the holding | • Number of courses used on the farm (calculated from quantity of each medicine used on farm and the quantity required per course)<br>• Number of cattle at risk of treatment, split into youngstock (<24months) and adult (>24months) | • Takes into account number of cattle and can use specific weights<br>• Takes into account course differences across AMs<br>• Takes into account youngstock use (and can separate use in youngstock from use in adults) | • Requires information on the number of youngstock as well as the number of adult dairy cattle<br>• If using standardised weights (100 kg and 600 kg) and/or defined dose rates and course lengths, these may not be accurate for a particular country or<br>unit<br>• Medicines are assigned only for youngstock or only for adults, which may not be accurate | T. Clarke, personal communication |

\* Rather than dose per kg liveweight, lactating cow tubes are dosed at number of tubes per cow per day, dry cow tubes are dosed as '4 per cow' as a single treatment and intrauterine products are one unit per cow.

### Comparison of metrics using real data

In order to further illustrate the different metrics, data on AM use for a 12-month period during 2015-2016 from six farms currently enrolled in local AMR research was collated (Figure 2). Here, the six farms were selected from a wider project to represent specific segments of the industry: Farms A and B are high-yielding, intensive, indoor Holstein herds; Farms C and D are extensive, block-calving, crossbred herds; Farms E and F are average-sized, average production Holstein-Friesian herds. Further details of the management practices and data collection on these farms can be found in Supplementary Material S1 and Table S1.

The AM use of these farms is represented by total mg/kg, DDDvet, DDDUK, DDDfarm, DCDvet, DCDUK and DCDfarm metrics. DDDvet and DCDvet were calculated using the ESVAC-defined weight of 425 kg using only adult stock numbers. DDDUK and DCDUK use dose rates and course lengths specific to the UK, obtained from SPCs and weights of 600 kg using only adult stock numbers. DDDfarm and DCDfarm use dose rates and course lengths specific to the farm and weights of 600 kg for adults and 100 kg for animals under 12-months of age (because for DDDfarm and DCDfarm, antimicrobials can be assigned to adults or calves). Total mg/kg also assigns antimicrobial use to adults or calves, with the same weights as DDDfarm.

A number of medicines were excluded across all metrics to allow comparability of benchmarking (e.g. where DDD values were not defined). Further details of the assumptions made are given in Supplementary Material S2; Figures S1 and S2 show the sensitivity of the metrics to these exclusions.

The illustrative metrics are presented for total AMs and for the HP-CIAs (3rd and 4th generation cephalosporins and fluoroquinolones) separately. In each panel, farms are ranked according to their use by that metric. It is interesting to note that the ranking is not the same across all metrics for total AM use (figures on the left) although it is largely the same for HP-CIA use (figures on the right).

Some interesting points are illustrated by these panels. For instance:

- Farm A is ranked last when using ESVAC or UK-level dose or course metrics for total use (DDDvet, DDDUK, DCDvet and DCDUK) but, using mg/kg and farm-specific metrics (DDDfarm and DCDfarm), it falls in the middle. This variation is primarily due to use of intramammary tubes with double daily doses and longer course lengths (beyond recommendations) under the Cascade. This increases dose and course metrics that assume standard doses have been given, but, given the relatively low amount of active ingredient in intramammary tubes, doesn’t significantly increase mg/kg. However, Farm A uses no HP-CIAs.
- Farm B and F swap rank when comparing DDDvet with DDDUK, and DCDvet with DCDUK for total use, indicating the difference between ESVAC and UK-level metrics.
- Farm C and D (extensive, block-calving, crossbred herds) are consistently low users across all metrics for total use, although Farm C uses more HP-CIAs.
- Farm E is ranked last when using farm-specific metrics for total use (DDDfarm and DCDfarm), but has a better position for ESVAC and UK course and dose-level metrics. This is because much of the injectable medicine used on this farm is used in calves and mg/kg, DDDfarm and DCDfarm split medicine use into adults and calves. These differences illustrate the point that more specific dosage and weight information can offer further insight into AMU on cattle farms, although this comes with the trade-off that more effort is required to gather more farm-specific data.
- Farm F is always benchmarked as the largest user of HP-CIAs but is never ranked as the largest in total use.

### Current use of metrics

Each metric presented in this manuscript is in common use. For example, in the UK, daily dose metrics are used by at least two retailers and CCC is currently in use by veterinary practices and retailers. Total mg/PCU is used to analyse UK-level sales data from pharmaceutical companies (17). DDDvet and DCDvet have been used in Ireland (12). Country-specific daily dose metrics are used in AMU reporting systems in the Netherlands (13) and in Denmark (via VetStat (11), Danish Integrated Antimicrobial Resistance Monitoring and Research Program (DANMAP) and the Danish Veterinary and Food Administration (DVFA)). Interestingly, DANMAP and the DVFA use different defined doses resulting in significant discrepancies in measurements of AM consumption using data from the same country (9). Daily dose metrics have also been used in studies outside of Europe (e.g. Canada (16) and Argentina (14)). Australia have published their overall antimicrobial figures as mg/PCU (25).

Current work at the University of Bristol with UK-based farmers, veterinarians and retailers as well as the experience in the Netherlands (26) suggests the need for specific metrics to be chosen and used consistently. These metrics need to be clearly explained so that users understand what data are required and the assumptions and biases behind the calculations. For a metric to be useful to farmers and veterinarians, it must be good for benchmarking purposes (i.e. it must be accurate and comparable at the unit level). This ideally means a metric that takes into account varying cattle numbers and weights as well as different management systems and does not penalise farmers or veterinarians for using medicines with higher mg/kg dose rates, such as the first-line AMs.

### Data issues and requirements

All metrics require accurate, representative and validated data and parameterisation in order to be useful. The section below discusses issues with obtaining detailed data and medicine information, and the assumptions and sensitivities around these data.

#### Data collection and data quality

Data for assessing AMU may come from the farmer or veterinarian (the actual usage amounts, cattle numbers and ages, average cattle weights, actual dosing and course regimens) and from regulatory bodies such as the EMA, Veterinary Medicines Directorate (VMD) or National Office of Animal Health (NOAH) (advised/defined dosing and course regimens for each medicine, mg of active substance per medicine unit). Cattle numbers may also be obtained from the BCMS in Great Britain and the Animal and Public Health Information System in Northern Ireland, which give information on all individual cattle on farms at any one time (N.B. as the reporting process is not perfect, these data are not always 100% accurate). Herd size can increase and decrease over an analysis period which may cause inaccuracies in the metric calculations if, for example, animal numbers are taken from one timepoint rather than using an average over the period.

While all farmers must keep records of medicine use in a medicine book (27), animal health and welfare tasks are likely to be prioritised instead, which can make record keeping a rushed exercise and lead to low quality data (28). Automating data entry on farms, with medicine recording linked to a standard identifier (e.g. VM number), could improve data quality. For example, teams such as VirtualVet (www.virtualvet.eu/) aim to develop systems allowing farmers to scan the bottle or pack of medicine, scan the eartag of the cattle being treated and add dose information. However, farm-side automation requires a robust system and hardware that is functional on farm as well as a concerted effort from farmers to use it. Automation on the side of the veterinarian is more straightforward as many veterinary practices already use practice management systems to enter sales data, and information can be extracted from these (as shown by VetIMPRESS, FarmVet Systems, www.vetimpress.com/), although this often requires substantial cleaning. However, the use of sales data assumes that all medicines sold to the farmer are used on that particular farm, for the animals specified, within the specified time period and at the correct dose rate, etc. (29). In fact, veterinarians anecdotally report that farmers may treat animals using unused medicines from previous sales; in these instances, it may be that the actual dose regimen does not match the recommended regimen for the medicine.

Some countries have strict monitoring systems for medicine sales in place, and similar methods could be implemented in the UK. The Netherlands, for example, requires veterinarians to upload sales data to “Medirund” within 14 days of the sale (www.medirund.nl/dierenarts/); this system then produces quarterly reports of AMU (using daily dose metrics) for both the veterinarian and the farmer.

#### Assumptions and sensitivities

In the absence of individualised weight data, the weight assumption used in calculations must be clearly stated and the sensitivity of the metrics to this assumption should be explored, particularly for comparison across farms or veterinarians. Similarly, assumptions and sensitivities about the treated population size, age and breed should be clearly presented in analyses.

If a metric is calculated for AM use over only a number of months (e.g. quarterly) there may be seasonal trends due to farm management (e.g. calving) that may skew the data; if instead a metric uses data from an entire year, these variations may be mitigated (although in some cases there may still be outliers). With all metrics, it is preferable to continually measure and monitor over time: more regular monitoring will provide more detailed information, aiding in understanding of the system and changes over time.

#### Medicine information

Every medicine licensed in animals in the UK has an SPC document which includes text summaries of recommended dose rates and course length for every animal in which that medicine is licensed for use, along with the active substances the medicine contains and their concentrations. In order to use these data in metric calculations, the values must be extracted from the text into an accessible format. Additionally, many SPCs present a range of doses or course durations, and a single value must be chosen, introducing potential inaccuracies (Table 4).

At present, the EMA and others do not calculate DDDvet and DCDvet at the product level, but rather at the level of the the active substance (i.e. oxytetracycline for each species and administration route). This means that there are currently no set standards for individually licensed products, and all medicines containing a certain active substance for use in a certain species will be using a single set dose rate or frequency. Work at the University of Bristol has also shown that there are some substantial differences in doses from UK SPCs to the doses accepted as part of the ESVAC DDDvet and DCDvet calculations. There are a number of medicines that have UK dose rates that are half or sometimes double those specified by ESVAC; these occur in the injectable antibiotics as well as oral preparations. To prevent duplication of effort and to standardise decisions on dose rates and frequencies appropriate to the UK, as well as to allow comparisons across analyses using these metrics, these authors recommend that a definitive list of standardised values for each of the licensed veterinary medicines in the UK is produced. This could be achieved by convening a workshop of key stakeholders to establish UK figures and including these values on future iterations of the downloadable Product Information Database Snapshot currently available on the VMD website (https://www.vmd.defra.gov.uk/ProductInformationDatabase/). Together with the VMD, the authors are currently producing such a list of medicines licensed in cattle in the UK to be published and maintained as a comprehensive standardised medicines database.

## Discussion

Many metrics have been presented and not one in itself is perfect. It is the assertion of these authors that the most elucidating metrics for the UK dairy industry would be UK-specific versions of the daily dose and course metrics using actual (or UK-estimated) cattle and youngstock weights along with actual (or defined UK-specific) treatment-level dose rates and course durations for medicines currently licensed in the UK. These UK-level metrics for livestock would also need clear assumptions for determining the number and type (specifically age) of animals at risk of treatment. Of course, a ‘gold-standard’ metric would use doses and courses specific to the unit (e.g. farm or veterinarian) and use actual animal number and weight data. This is a possible target for the future, as systems to collect farm-level, individual cattle data are being developed but are not currently in wide use (VetIMPRESS, VirtualVet).

If the UK could publish its standard daily doses for veterinary medicines in the same way as it does for human medicine (i.e. for every individual product rather than by active substance and route as are currently provided by ESVAC (23)), a UK-specific daily dose metric would be feasible and would also allow comparison of usage with the medical profession on a country-wide basis. Part of the required standardisation will be formalising the choice of medicines to include in the metric (and, indeed, to include in reduction targets). For example, how antibiotic sprays, footbaths and dry cow tubes should be included in daily dose metrics, and whether HP-CIAs should also be reported separately to total use and have additional targets.

Given the work required to tailor dose and course metrics for the UK, this may prove infeasible, at least in the short-term. As an alternative, the next best option may be the DDDvet and DCDvet metrics currently being standardised by ESVAC (10). Although these metrics provide Europe-wide generalisations of dose rates and course lengths and have the limitation of defining only by active substance, the availability of the standardisation makes these metrics appealing until such data can be generated for the UK.

As an alternative to daily dose metrics, total mg/kg is simple to calculate and understand and requires none of the standardisation decisions. If presented with the relevant caveats and presented separately for HP-CIA and non-HP-CIA medicines, mg/kg is suitable for tracking usage on a single unit (farm, veterinary practice or retailer) over time. However, mg/kg is less suitable for cross-unit comparisons (unless farm-specific weights are used).

This manuscript seeks to elucidate the pros and cons of the current metrics being considered for measuring AMU in the UK cattle industries. The ultimate reason for use of these metrics should be to aid efforts to reduce AMR and to encourage best practice stewardship of AMs across the livestock industries. It is recognised that encouraging low AMU needs to be balanced against maintaining animal health and welfare and that individual farm context is important to inform the most responsible use of AMs on that farm. In recent years, the move towards routine recording and collection of usage data at farm and veterinary practice level has steadily increased, and, although there are still questions about the quality of these types of data, improvements are being made continually as technology and software develop. Availability of reliable data would encourage parties to make use of these data to drive change. In the experience of these authors, farmers, veterinarians and retailers are all keen to use data to understand and reduce usage of AMs. The UK stands to learn much from other countries where such practices have already been employed and, with the current drive from VMD and others, this is becoming a priority.

## Acknowledgements

The authors are grateful to Tom Clarke from Synergy Farm Health for providing information about Cow Calculated Courses and to members of the Bristol Veterinary School’s Farm Animal Group for reviewing the manuscript.

